# Overflooding Ratios in the Sterile Insect Technique: Toward Sustainable Management of *Bagrada hilaris*

**DOI:** 10.1101/2025.03.05.641589

**Authors:** Chiara Elvira Mainardi, Chiara Peccerillo, Alessandra Paolini, René F. H. Sforza, Alessia Cemmi, Ilaria Di Sarcina, Francesca Marini, Massimo Cristofaro

## Abstract

The Sterile Insect Technique (SIT) is an eco-friendly control method that may prove effective against *Bagrada hilaris*, a pest native to India, Southeast Asia, and Middle and central Africa and reported as invasive in the southwestern USA, Hawaii, Mexico, South America, and two Mediterranean islands. This insect causes significant crop damage due to its intense feeding behavior and is currently managed almost exclusively with synthetic insecticides. In this context, SIT offers a promising alternative for controlling *B. hilaris* populations, provided that sufficient numbers of sterile males are continuously released. Based on this premise, we conducted a preliminary laboratory study to evaluate the overflooding ratio (OFR)—the proportion of non-irradiated to irradiated males required to suppress the population’s fertility. We tested various OFRs (1:1, 1:2, 1:5, and 1:10), monitoring both the number of eggs laid and hatching rates. Our results show a significant decrease in fertility as the percentage of irradiated males increases. Among the ratios examined, 1:5 emerged as the most advantageous in terms of both reducing fertility and ease of application. Although further validation under field conditions is needed, our findings suggest that SIT could effectively contribute to an integrated management strategy for *B. hilaris*, reducing the reliance on chemical pesticides and supporting a more sustainable approach.

## 1. Introduction

The Sterile Insect Technique (SIT), conceptualized by Knipling in the 1930, is an autocidal pest control method [1] involving the mass rearing of target insects, sterilization of males through gamma or X-ray irradiation, and their periodic release into the environment [2]. Wild females that mate with these treated males produce sterile eggs, resulting in a gradual reduction of the target population [3]. SIT is considered an ecological approach, as it does not affect non-target organisms and is not subject to resistance phenomena [3], which has led to its increasing adoption worldwide [4].

This technique is particularly straightforward to apply to holometabolous insects, due to their quiescent pupal stage that facilitates operations [5]. By contrast, one requirement for applying SIT to a target species is that adults must not be the principal cause of damage [5]. In hemimetabolous insects, if adults cause significant crop damage, releasing large numbers of sterile individuals could lead to immediate losses [5]. Nevertheless, in restricted areas such as greenhouses and isolated cropping systems, SIT can still contain the pest and ultimately achieve population control [6]. For these reasons, it has been proposed as a control strategy for the invasive species *Bagrada hilaris* [7].

The painted bug, *Bagrada hilaris* (Burmeister) (Hemiptera:Pentatomidae) is currently regarded as a serious economic pest of several vegetable crops, mainly belonging to the Brassicaceae family [8]. Originally distributed in Africa, The Middle East, and Southern Asia [9,10], it has recently been reported in California, central Arizona [8], Nevada, New Mexico, and Utah [11].Recently, it has been recorded in Mexico [12], Hawaii [13], Chile [14] and Argentina [15]. In Europe, *B. hilaris* was first identified in the late 1970s on the island of Pantelleria (Italy) and Malta [16]; since then, it has established mainly on caper plants (*Capparis spinosa* L.), becoming a key pest of this crop in the island [17].

*Bagrada hilaris* is distinguished from over 4,700 other Pentatomid species [18] by its peculiar oviposition behavior [13,19]. Specifically, the female lays single eggs or small clusters [9,20,21] under a layer of sand [21,22].

This pest attacks crops belonging to at least 23 different families [13], with particularly severe economic impacts on Brassicaceae [23] and, on the island of Pantelleria, on capers [17,24]. Feeding by *B. hilaris* causes chlorotic lesions that may become necrotic and either reduce plant growth or lead to plant death [13,20,23,24]. Moreover, its gregarious behavior [25] amplifies the damage, and in some parts of the USA and Mexico, agricultural losses have exceeded one billion U.S. dollars [8,26].

Common control practices mainly rely on repeated applications of broad-spectrum insecticides (particularly pyrethroids and carbamates) [13,26]. However, many growers in the USA prefer to avoid synthetic pesticides and instead modify their cropping systems to reduce the economic losses of *B. hilaris* [20]. The lack of a classical biological control program in newly invaded areas [27] further underscores the need for more sustainable strategies. In this context, SIT represents a valid alternative. Since *B. hilaris* females cause the most severe damage [23], releasing only sterile males could limit the overall harm, especially if combined with other biological control methods [28]. Given these considerations, the island of Pantelleria (Italy) is an ideal model area where SIT could effectively control the population of *B. hilaris*, owing to its insular geographic characteristics.

Recent studies have examined the effects of gamma irradiation on adult *B. hilaris*, showing that sterilization rates of 90% are achieved at 60 Gy, 95% at 80 Gy, and total suppression of fertility at 100 Gy absorbed dose and above [7]. The 80 Gy dose was found to be optimal, ensuring a high level of sterilization while maintaining adequate sexual performance. Indeed, female *B. hilaris*—both under choice and no-choice conditions—favors mating with irradiated males, which spend more time copulating than unirradiated males [29]. The sterile eggs resulting from irradiated male × fertile female matings (SIT eggs) can be integrated into classical biological control programs, using specific egg parasitoid wasps, such as *Gryon aetherium* Talamas (Hymenoptera: Scelionidae), which successfully parasitize these eggs [30].

The synergy between SIT and existing biological control approaches [28,31] strengthens the possibility of containing *B. hilaris* in infested areas [6,32].

The effectiveness of SIT depends on the size of the target population, the extent of the treated area, the program’s objectives [33], and the required ratio of sterile to wild insects in the field. This ratio, known as the overflooding ratio (OFR), precisely estimates the optimal proportion of sterile to wild males. If the number of released sterile males does not substantially exceed that of native males, the induced sterility will be inadequate [5]. In essence, the induced level of sterility must overcome the reproductive rate (i.e., the mating success) of wild females [34]. Several mathematical models have been developed to determine the minimum OFR required to suppress target populations. Knipling [3,35] introduced a simplified geometric population growth model, later expanded by Berryman [36] to incorporate a competitiveness factor (which compares the effectiveness of sterile vs. wild males). Finally, Barclay [37] refined the model to include population growth rate, survival, dispersion, release frequency, and economic/logistical considerations. While these models can offer high theoretical precision, they often require parameter estimates (e.g., male/female population size, intrinsic growth rates) that are difficult to measure under field conditions. Several studies have provided OFR estimates for different species: around 20:1 for *Ceratitis capitata* (Wiedemann) (Diptera: Tephritidae)[38,39], *Bactrocera dorsalis* (Hendel) and *Bactrocera cucurbitae* (Coquillett) (Diptera: Tephritidae) [39], and species from the genus *Anastrepha* such as *A. ludens* (Loew) and *A. obliqua* (McQuart)(Diptera: Tephritidae) [40].

Since the OFR is influenced by seasonal variables, it must be viewed as a dynamic parameter. Pest populations do not continuously increase at high rates; rather, they reproduce in cycles related to biotic and abiotic factors such as food availability, weather conditions (temperature, humidity), and crop cycles [41]. Hence, integrating these factors into monitoring and release planning is crucial to ensure that the OFR remains effective in reducing the target population. Despite the advances in understanding the irradiation effects on *B. hilaris*, no studies have yet quantified the OFR required to curtail its reproductive capacity.

For this reason, the present work aims to conduct a preliminary laboratory study to assess how different OFRs may efficiently induce sterility. The resulting data are discussed in the context of releasing *B. hilaris* individuals on Pantelleria Island (located South of Sicily, Italy) in combination with classical biological control, ultimately seeking to prevent any further expansion of this species in the region.

## 2. Materials and Methods

### 2.1. Insect collection and rearing

During late summer and autumn 2023, individuals of *B. hilaris* were collected on Pantelleria Island, in the Scauri area (36°46’23.0”N 11°57’41.0”E), at the “Cooperativa Agricola Produttori Capperi”. The number of specimens collected varied by season, ranging from one hundred to one thousand individuals per sampling event.

Collection was facilitated by an entomological aspirator, taking advantage of the insects’ aggregative behavior, and included both adults and nymphs of various instars from caper plants (*Capparis spinosa* L.). The capture was most successful during the hottest hours (late morning to mid-afternoon). Collected specimens were placed in cardboard tubes (4.1 × 11.9 cm) and transported to the quarantine facilities of the Edmund Mach Foundation in San Michele all’Adige, Trento, Italy. There, insects were transferred into BugDorm® cloth cages (30 × 30 × 30 cm, 680-μm mesh), maintained at 25 ± 1°C with a 14:10 h (L:D) photoperiod and 60% relative humidity. Brussels sprouts (*Brassica oleracea* L. var. *gemmifera*) were provided as a food source, replaced three times per week. A plastic Petri dish (9 cm Ø) filled with fine sand (particle size ≈200 µm) was placed in each cage as an oviposition substrate. Field-collected insects were periodically added to the colony to maintain genetic diversity.

### 2.2 Irradiation

Fifth-instar nymphs were selected from the laboratory colony and placed in 50 mL CorningTM Falcon plastic tubes. These individuals were then isolated in Petri dishes (5 cm Ø) until they reached adulthood. This procedure allowed the separation of males and females based on sexual dimorphism [13] and ensured their virginity. Groups of ten males were then placed in Petri dishes (9 cm Ø) and shipped from the Edmund Mach Foundation to Rome (Italy) via DHL using “5-level” packaging to ensure safe transport.

The insects were irradiated at a dose of 80 Gy at the Calliope gamma irradiation facility [42] at the ENEA Casaccia Research Center (Rome). Calliope is a pool type irradiation facility equipped with 25 Cobalt-60 radioisotope sources (average energy 1.25 MeV). Irradiation was performed at an average dose rate of approximately 160 Gy/h (2.67 Gy/min). The dose rates were determined by Fricke dosimetry, and the corresponding values are referenced to water.

The irradiation dose was chosen because it has been indicated as optimal for SIT application [29].

Non irradiated males and females, intended for use as controls, were also transported to the same irradiation facility. At the end of the process, all specimens were returned to the Edmund Mach Foundation quarantine facility, following the same procedure.

### 2.3 Experimental Setup

The experiment was started the day after irradiation. BugDorm® cloth cages (17.5 × 17.5 × 17.5 cm, 160-μm mesh) each received a small Petri dish (5 cm Ø) filled with fine sand and, depending on the total number of individuals, two or three Brussels sprouts were added as food source. In each cage, one non-irradiated virgin female was confined together with non-irradiated and irradiated males in the following ratios: 1:1:1, 1:1:2, 1:1:5, and 1:1:10 (one female, non-irradiated males:irradiated males). The control group consisted of one healthy virgin female confined with 2, 3, 6, or 11 non irradiated males. In this way, the females were confined to the same number of treatment males in each relative control.

The temperature, humidity and L:D conditions were the same as in the rearing. Every five days, the food was replaced, and the Petri dish was removed and then replaced with a new one with clean (=untouched) sand. Each Petri dish containing sand was inspected under a stereomicroscope (Olympus SZX-ILLB200, Hamburg, Germany) to count the number of eggs laid. Subsequently, the Petri dishes with eggs were maintained under the same environmental conditions to evaluate hatching rates. Starting five days after the initial count, hatched nymphs were recorded. The following hatching checks occurred every 5 days up to a maximum of 20 days. The eggs that did not hatch within 15 days were deemed non-viable, and hatching checks ceased. Different replicates were observed until the death of at least one male or the female. For each replicate, four observations were performed over a total of 20 days. The observations were grouped in 4 categories corresponding to the days of the check starting five days after the beginning of the experiment (Observation 1 = check occurred on the fifth day, Observation 2 = check occurred on the tenth day, Observation 3 = check occurred on the fifteenth day, Observation 4 = check occurred on the twentieth day). Finally, the effect of the different ratios on fertility was assessed, expressed as the ratio of the number of hatched nymphs to the total number of eggs laid.

### 2.4 Statistical analysis

A non-linear least squares algorithm was used to estimate fertility, defined as the ratio between the number of nymphs and eggs, based on the number of irradiated males. This approach allowed us to compute the expected frequency based on a hypothetical number of irradiated males. Model estimates predictions were calculated using the functions nls [43] and predict NLS. Fertility differences among groups were assessed using the Kruskal-Wallis test [43], followed by post-hoc comparisons using Dunn test with Bonferroni correction. A non-parametric test was chosen because fertility did not follow a normal distribution in each group, as indicated by the Shapiro test.

All analyses were conducted using a significance level of α = 0.05. Statistical analyses were performed using R (version 4.3.1). R packages readxl [44], ggplot2 [44],flextable, gtsummary [45] and dplyr [46] were used to import, manipulate and export results.

## 3. Results

### 3.1 Anova test on the fertility value in the groups given by the ratio

**Figures** from **1** to **4** show fertility trends (% eggs hatched) in *B. hilaris* at four different ratios of unirradiated males: irradiated males (1:1, 1:2, 1:5, 1:10), compared with the correspondent control group consisting of only unirradiated males. **Figure 1** shows no significant difference between control and treatment in any observation group. In **Figure 2**, there is a significant difference in Observation 4 (p=0.0319). In **Figure 3**, only Observation 1 shows a significant difference between treatment and control (p=0.0057). In the last graph (**Fig. 4**), there were significant differences in Observation 2 (p=0.0003) and Observation 3 (p=0.0349).

**Figure 1:**
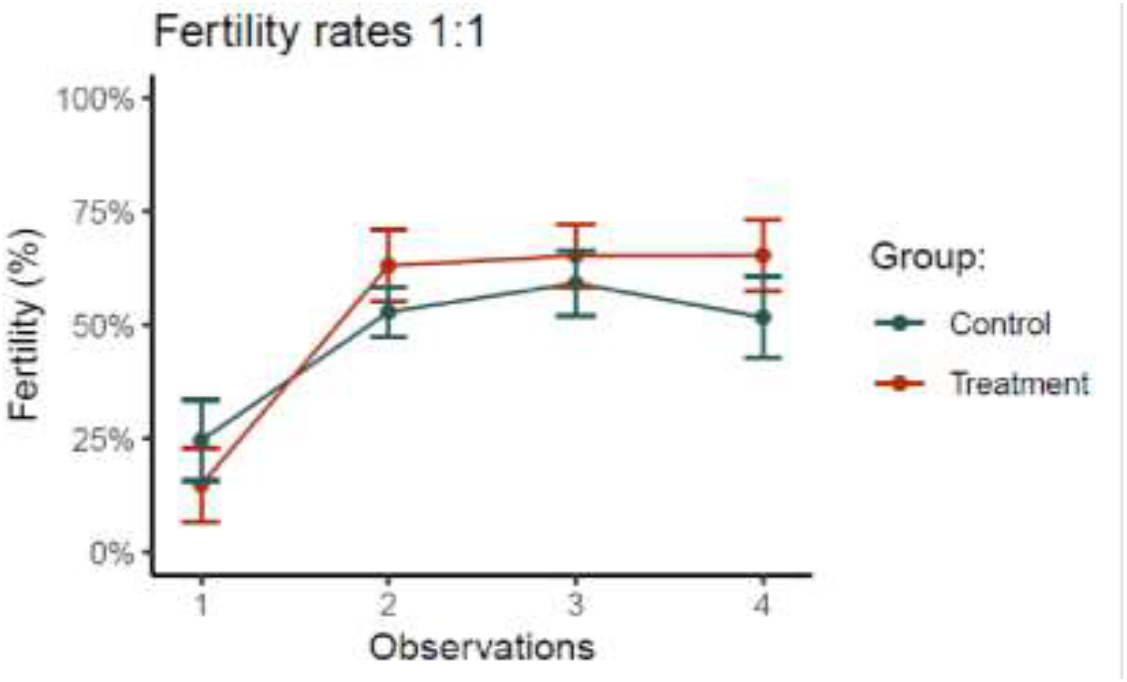
Broken-line graph showing the fertility (%) of *Bagrada hilaris* females under two different mating conditions across four time points (Observations). The red line (treatment) represents one female confined with one irradiated male and one non irradiated male (1:1:1). The green line (control) represents one female confined with two non irradiated males (1:2). Observations 1, 2, 3, and 4 correspond to days 5, 10, 15, and 20, respectively, from the start of the experiment. Vertical bars depict standard errors.

**Figure 2:**
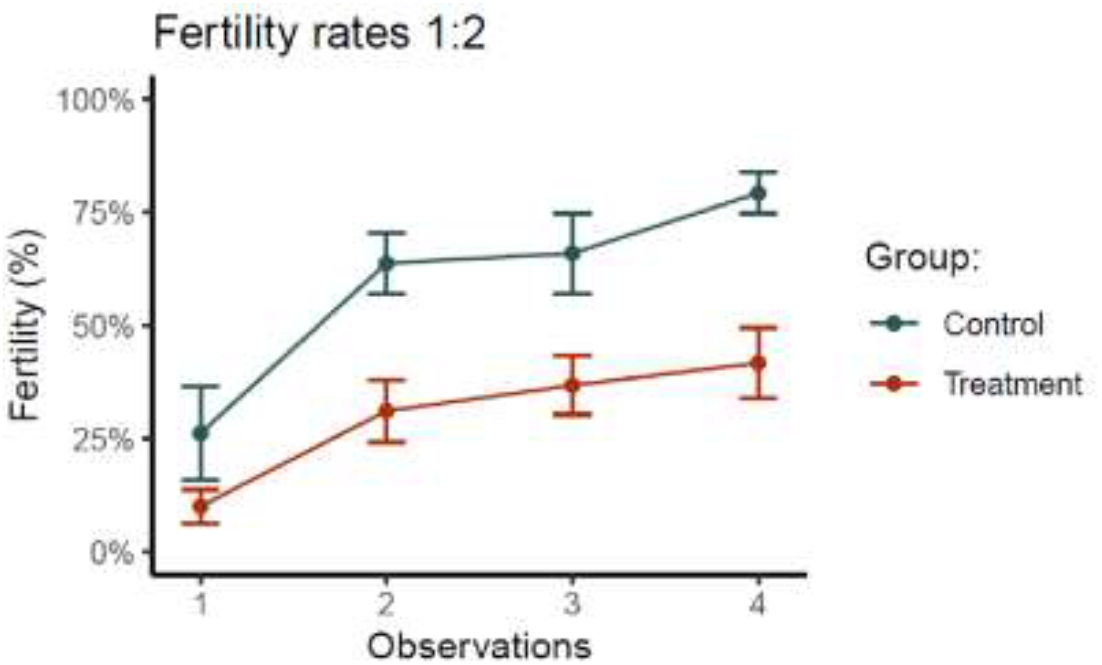
Broken-line graph showing the fertility (%) of *Bagrada hilaris* females under two different mating conditions across four time points (Observations). The red line (treatment) represents one female confined to one non irradiated male and two irradiated males (1:1:2). The green line (control) represents one female confined to three non irradiated males (1:3). Observations 1, 2, 3, and 4 correspond to days 5, 10, 15, and 20, respectively, from the start of the experiment. Vertical bars depict standard errors.

**Figure 3:**
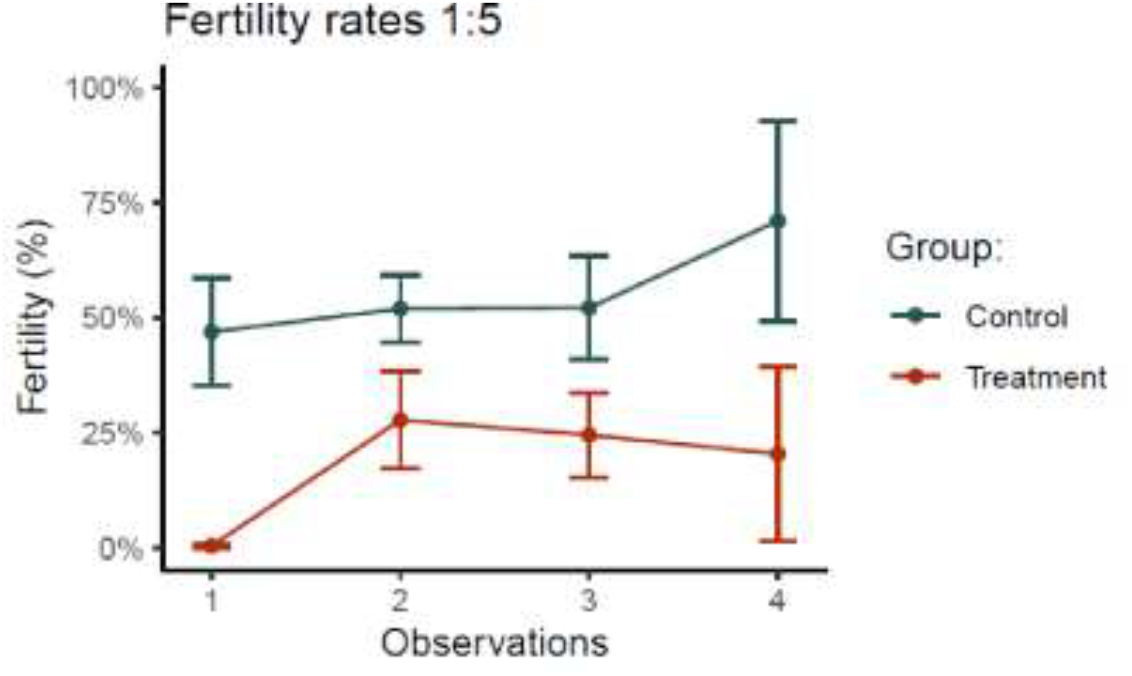
Broken-line graph showing the fertility (%) of *Bagrada hilaris* females under two different mating conditions across four time points (Observations). The red line (treatment) represents one female confined to one non irradiated male and five irradiated males (1:1:5). The green line (control) represents one female confined to six non irradiated males (1:6). Observations 1, 2, 3, and 4 correspond to days 5, 10, 15, and 20, respectively, from the start of the experiment. Vertical bars depict standard errors.

**Figure 4:**
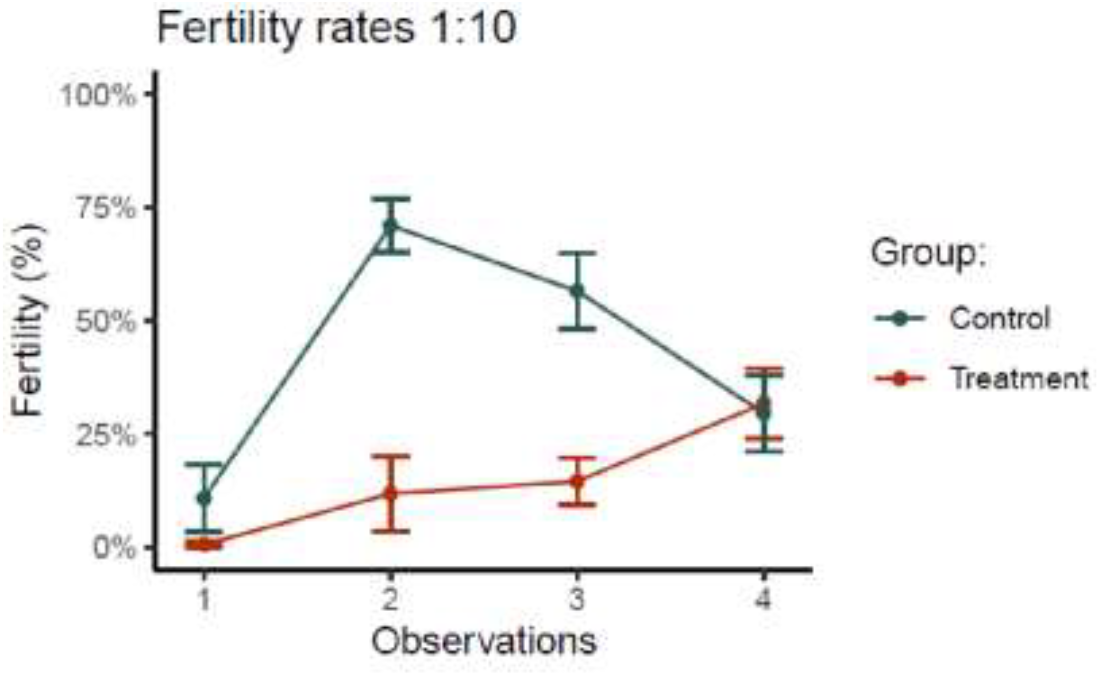
Broken-line graph showing the fertility (%) of *Bagrada hilaris* females under two different mating conditions across four time points (Observations). The red line (treatment) represents one female confined to one non irradiated male and ten irradiated males (1:1:10). The green line (control) represents one female confined to eleven non irradiated males (1:11). Observations 1, 2, 3, and 4 correspond to days 5, 10, 15, and 20, respectively, from the start of the experiment. Vertical bars depict standard errors.

Statistical analysis that takes a comprehensive look at all treatments (**Tab. 1**) shows that for the Observation 1 group there is a significant difference between the ratio 01:06:00 and 01:01:05 (p-value=0.0076), and between 01:06:00 and 01:01:10 (p-value=0.0057).

**Table 1:** Kruskal Wallis test comparison between treatment and control groups per observation.

| Observation | Ratio | Number of observations | Average fertility | Kruskall-Walli's test statistics (df) | p-value |
| --- | --- | --- | --- | --- | --- |
| **1** | 01:01:0<br>1 | 16 | 14.92 | 19.131<br>(7) | 0.008 |
|  | 01:01:0<br>2 | 21 | 10.16 |  |  |
|  | 01:01:0<br>5 | 12 | 0.38 |  |  |
|  | 01:01:1<br>0 | 13 | 0.64 |  |  |
|  | 01:02:0<br>0 | 14 | 24.70 |  |  |
|  | 01:03:0<br>0 | 14 | 26.27 |  |  |
|  | 01:06:0<br>0 | 10 | 46.93 |  |  |
|  | 01:11:0<br>0 | 12 | 10.88 |  |  |
| **2** | 01:01:0<br>1 | 16 | 63.19 | 34.602<br>(7) | <0.001 |
|  | 01:01:0<br>2 | 21 | 31.20 |  |  |
|  | 01:01:0<br>5 | 12 | 27.74 |  |  |
|  | 01:01:1<br>0 | 13 | 11.90 |  |  |
|  | 01:02:0<br>0 | 14 | 52.96 |  |  |
|  | 01:03:0<br>0 | 14 | 63.81 |  |  |
|  | 01:06:0<br>0 | 10 | 51.94 |  |  |
|  | 01:11:0<br>0 | 12 | 70.99 |  |  |
| **3** | 01:01:0<br>1 | 16 | 65.36 | 31.96<br>(7) | <0.001 |
|  | 01:01:0<br>2 | 21 | 36.87 |  |  |
|  | 01:01:0<br>5 | 12 | 24.49 |  |  |
|  | 01:01:1<br>0 | 13 | 14.61 |  |  |
|  | 01:02:0<br>0 | 14 | 59.31 |  |  |
|  | 01:03:0<br>0 | 14 | 66.01 |  |  |
|  | 01:06:0<br>0 | 10 | 52.15 |  |  |
|  | 01:11:0<br>0 | 12 | 56.58 |  |  |
| **4** | 01:01:0<br>1 | 13 | 65.46 | 23.406<br>(7) | 0.001 |
|  | 01:01:0<br>2 | 13 | 41.80 |  |  |
|  | 01:01:0<br>5 | 4 | 20.37 |  |  |
|  | 01:01:1<br>0 | 6 | 31.76 |  |  |
|  | 01:02:0<br>0 | 12 | 51.81 |  |  |
|  | 01:03:0<br>0 | 11 | 79.30 |  |  |
|  | 01:06:0<br>0 | 3 | 71.06 |  |  |
|  | 01:11:0<br>0 | 5 | 29.67 |  |  |

Within the Observation 2 group, there is a significant difference between the ratio 01:01:10 and 01:01:01 (p-value=0.0009), 01:03:00 and 01:01:10 (p-value=0.0014), 01:11:00 and 01:01:02 (p-value=0.0205), 01:11:00 and 01:01:05 (p-value=0.0203), and between 01:11:00 and 01:01:10 (p-value=0.0003).

For the Observation 3 group, the significant differences are between the ratio 01:01:05 and the ratio 01:01:01 (p-value=0.0168), 01:01:10 and 01:01:01 (p-value=0.0005), 01:02:00 and 01:01:10 (p-value=0.0099), 01:03:00 and 01:01:05 (p-value=0.0007), 01:03:00 and 01:01:10 (p-value=0.0007), and 01:11:00 and 01:01:10 (p-value=0.0349).

Within the Observation 4 group, the pairs of ratios with a significant difference are 01:03:00 and 01:01:10 (p-value=0.0163), 01:03:00 and 01:011:02 (p-value=0.0319), 01:03:00 and 01:01:05 (p-value=0.0407), and 01:11:00 and 01:03:00 (p-value=0.0203).

### 3.2 Model for estimating fertility based on the number of irradiated males

A non-linear least squares algorithm was used to estimate fertility, i.e. the ratio between the number of neanids and eggs (**Fig. 5**), based on the number of irradiated males.

**Figure 5:**
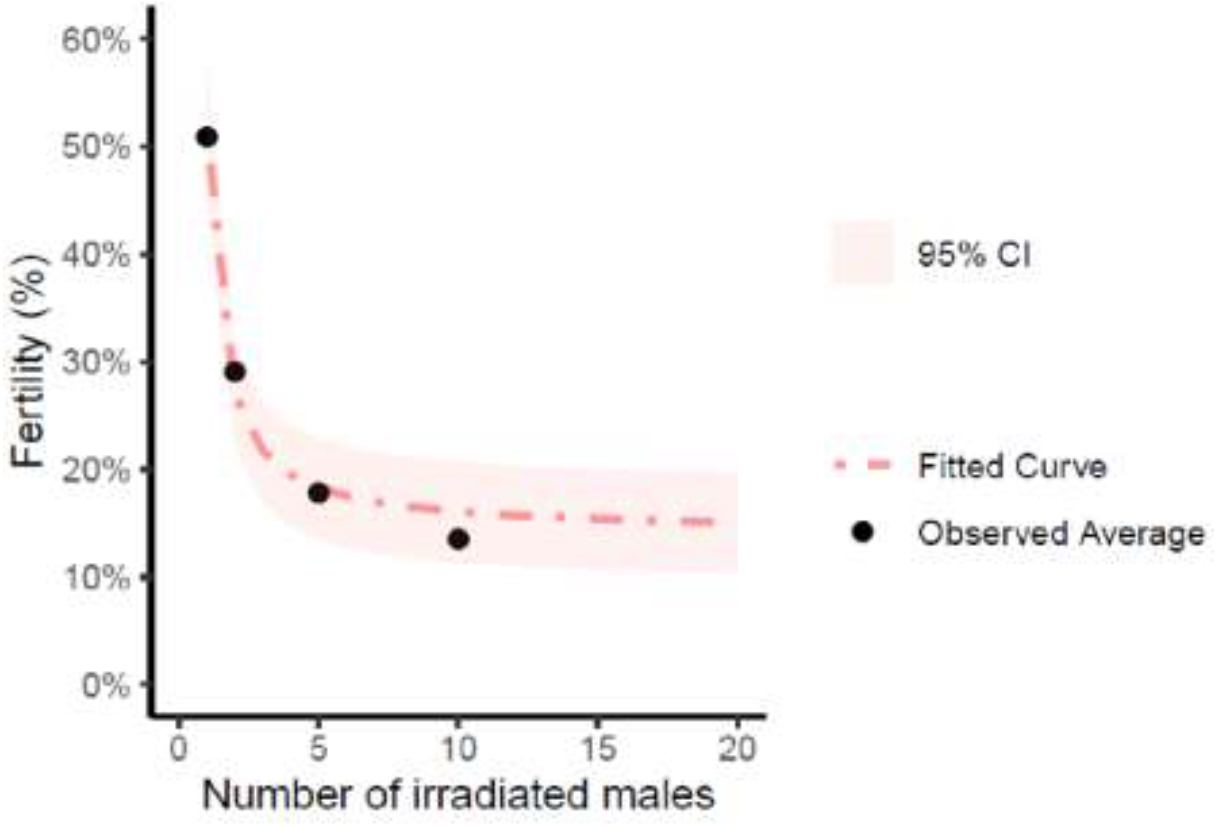
Fitted curve of the relationship between variable Fertility and number of irradiated males.

This approach is justified by observing the distribution of the average fertility value observed for the 4 scenarios tested.

An exponential equation was hypothesized by observing the distribution of average fertility by the number of irradiated males.

In particular, the hypothesized equation is of the type:

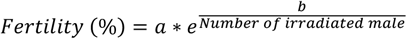

The nls function of the R software was used to estimate the parameters a and b of the equation. Parameter “a” represents a multiplicative factor, while “b” represents the speed of decreasing of the rate (**Tab.2**).

**Table 2:** Equation parameters estimated using NLS function.

| Parameters | Estimate | 95% CI<sup>1</sup> | p-value |
| --- | --- | --- | --- |
| a | 14.14 | 9.5, 19 | <0.001 |
| b | 1.29 | 0.91, 1.7 | <0.001 |

<sup>1</sup>CI = Confidence Interval

For the estimation of the parameters, all 252 observations for which the number of irradiated males was not missing were used (i.e. the observations referring to tested scenarios).

The model estimates the parameter a to be 14.14 and the parameter b to be 1.29. Both estimates are significantly different from 0 (p<0.05).

Table 3 presents the expected frequency based on the number of irradiated males considered. Specifically, when 10 irradiated males are used, the estimated reduction in fertility is 16.09%. This effect decreases as the number of irradiated males increases, reaching 15.08% at the maximum considered number of 20.

**Table 3.**
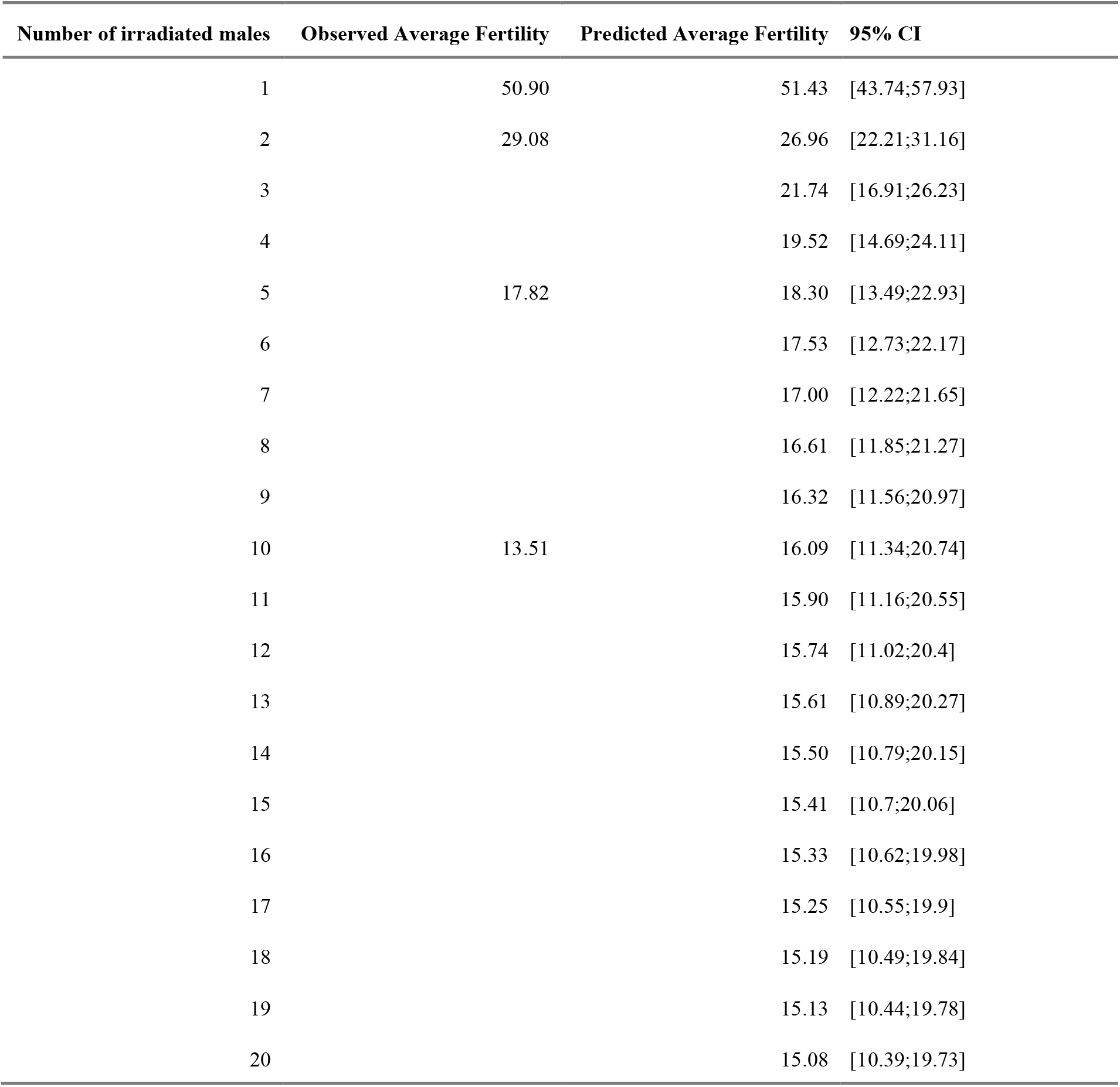
Observed and Predicted Average Fertility based on number of irradiated males. Predictions and confidence intervals are calculated using first order Taylor expansions.

## 4. Discussion

Successful application of the sterile insect technique occurs only when a sufficiently constant and effective inundative release of sterile individuals is ensured [38]. In the current study, our aim was to compare the levels of induced sterility in *Bagrada hilaris* males across different overflooding ratios.

Our results confirm that increasing the ratio of irradiated to non-irradiated males leads to a significant reduction in fertility. Specifically, the analyses revealed substantial differences among the four tested ratios (1:1, 1:2, 1:5, 1:10), with a marked decrease in the percentage of fertile eggs as the number of irradiated males rose. This trend is well described by a nonlinear exponential model, showing how fertility declines as the proportion of irradiated males increases.

In almost all overflooding ratios, the fertility of the treatment groups remained consistently lower than that of the controls. At the 1:1 ratio, both the control and treatment groups started at low fertility levels before experiencing a clear increase. In the end, the treatment reached slightly higher values than the control, stabilizing between 60–70% in both groups. At the 1:2 ratio, fertility began around 10% and increased over time, albeit without reaching control levels. The difference became far more significant for the 1:5 and 1:10 ratios: the treatment started with fertility values close to 0% and, although rising gradually, remained below 30%. Meanwhile, controls could exceed 70– 80% hatching by the final observations, revealing a wide gap.

Model analysis indicates that as the number of irradiated males increases (from 1 to 20), predicted fertility declines sharply and aligns closely with the values actually recorded, suggesting a good match between the model and the experimental data. From 10 irradiated males onward, predicted fertility drops by only 1–2 percentage points (from 16.09% to 15.08%), indicating that further increases in the number of irradiated males lead to only modest additional fertility reductions, at the cost of higher logistical and economic investment.

Overall, both the 1:5 and 1:10 ratios achieve a substantial reduction in fertility (about 17% vs. 13%, respectively). However, the difference between them does not appear sufficient to justify doubling the number of irradiated males. Consequently, the 1:5 ratio may be more favorable from an operational standpoint, ensuring a significant reduction in fertility while requiring fewer resources for rearing, irradiation, and release. Indeed, within the SIT framework, the goal is to identify an ideal overflooding ratio that ensures a strong impact on the pest population while remaining sustainable in the long term. Assuming our cage-based results can be reasonably extrapolated to open-field conditions, an OFR of at least 1:5 could reduce populations by up to 80%. At present, no reference values exist for other Pentatomids, except for *Halyomorpha halys* Stål (Hemiptera:Pentatomidae), where an OFR of 1:5 led to a 52% reduction in hatching, compared to an expected 62% [6].

It is important to note that our experimental conditions might underestimate the OFR needed: cage confinement could confer certain advantages to irradiated males by eliminating the need for long-distance dispersal to locate wild females. Consequently, higher OFRs might be required in the field. Additionally, various ecological and behavioral factors can alter the ratio of irradiated to non-irradiated males. For instance, the pest’s heterogeneous spatial distribution [47,48] and the need for irradiated males to locate the same aggregation sites as wild males [49] may demand further releases. Resource availability also affects local density [5], while in the case of *B. hilaris*, although polyphagous, it predominantly concentrates on caper in Pantelleria, potentially simplifying SIT deployment. On the island, caper fields are small agricultural units of couple hundred square meters, isolated or merged together, but separated by rock stone fences. This unique agricultural setting may be favourable for SIT as released irradiated males would easily remain close to the point of release. Moreover, the competitiveness of irradiated males is critical [50]: in *B. hilaris*, these males maintain reproductive performance comparable to wild males [51,52], reducing the need for additional releases. Finally, the presence of natural predators can lower the threshold of sterile males required [53]. There is no information on the natural predators of Bagrada on the island of Pantelleria, but spiders, mantids, and predatory Heteroptera (*Zelus* spp.) have been observed in the United States, feeding on neanids and adults [13].

In many instances, SIT is integrated with other control methods, as it proves more effective at low pest densities [37]. Indeed, it is common practice to reduce the wild population with insecticides or other control methods prior to releases. In the case of *B. hilaris*, for example, the egg parasitoid *Gryon aetherium* Talamas is capable of parasitizing even SIT eggs (fertile female × irradiated male) [54], suggesting a possible synergy between SIT and classical biological control [30,55].

Another important factor is the longevity of irradiated males: in multivoltine species, overlapping generations require continuous releases. The optimal release frequency (weekly, biweekly, or even daily) depends on how long sterile males survive in the field [56]. Moreover, releases must be carried out when the pest population reaches its seasonal minimum [33,34,57] to maximize the chances of success. For *B. hilaris*, adults emerge between late May and early June [17] and persist until February of the next year (personal observations). This timeline makes it possible to identify a natural decline phase—around March and April—during which releases could be initiated.

To more accurately define the OFR, a specific model must be developed that incorporates population size, growth rate, and residual fertility of irradiated males [37]. For *B. hilaris*, population size can be approximated through trap capture data, growth rate derived from egg-laying frequency, and residual fertility already established (95% sterilization at 80 Gy). The next step will be to test the proposed ratio (1:5) in situ, through semi-field trials, in order to assess its actual effectiveness under natural conditions.

In addition, SIT–CBC synergy was primarily examined in the context of the United States, where *B. hilaris* shows a haplotype closely linked to the Pakistani one [58]; moreover, *Gryon aetherium* has been reported in specific American regions (e.g., California and Chile; [59,60]). Hence, the ideal strategy in that areas would be to increase the adventive parasitoid population while providing ecological conditions that enhance its stabilization. Conversely, on Pantelleria Island, the Bagrada haplotype is more similar to the Maltese and Moroccan variants (Sforza, R.F.H., personal communication). Consequently, conspecific parasitoids may be present, and further investigations in the pest’s native range are needed, for instance via sterile sentinel eggs [54].

An alternative strategy involves the use of the “Kamikaze Wasp Technique” (KWT) [61], which innovatively combines CBC and SIT in a control operation while avoiding effects on non-target organisms. The pest can be controlled by rearing, sterilizing, and releasing both the target host and candidate parasitoid. The sterility of the parasitoid, in fact, makes it “single-use,” because the releases will not result in a self-sustaining population. It should be noted that some parasitoids require the input of venom and other factors injected by the mother into the host for development, as the developing parasitoid does not kill the host [61]. Thus, host development is blocked by the development of the parasitoid, which simultaneously inhibits its own progeny as a result of irradiation [61]. This strategy opens new perspectives for studying the effects of irradiation on the biological control candidate agents of *B. hilaris, Gryon aetherium*.

In conclusion, our data suggest that an OFR of at least 1:5 exerts a significant impact on the fertility of *B. hilaris*, offering a promising option for SIT applications. This study represents an initial step toward practical development of the technique for *B. hilaris*, while encouraging further field data collection. The synergistic effect of SIT alongside classical biological control may further bolster *B. hilaris* management, curbing insecticide use and promoting a more sustainable approach.

## Acknowledgments

The authors are grateful to all colleagues and friends who contributed to the study in various ways: F. Di Cristina, S. Barlattani, M. Guedj (BBCA onlus, Rome, Italy), for technical support; Gerardo Roselli and Gianfranco Anfora (Edmund Mach Foundation) for scientific contribution; R. Cappadona (Cooperativa Capperi Pantelleria, Italy), S. Anelli, A. Biddittu (Parco Nazionale di Pantelleria, Italy), G. Lasagni, and F. Maggiore (Bonomo & Giglio srl) for facilitating the field activities in the Island of Pantelleria.

